# Cross-linking BioThings APIs through JSON-LD to facilitate knowledge exploration

**DOI:** 10.1101/167353

**Authors:** Jiwen Xin, Cyrus Afrasiabi, Sebastien Lelong, Julee Adesara, Ginger Tsueng, Andrew I. Su, Chunlei Wu

## Abstract

**Background:** Application Programming Interfaces (APIs) are now widely used to distribute biological data. And many popular biological APIs developed by many different research teams have adopted Javascript Object Notation (JSON) as their primary data format. While usage of a common data format offers significant advantages, that alone is not sufficient for rich integrative queries across APIs.

**Results:** Here, we have implemented JSON for Linking Data (JSON-LD) technology on the BioThings APIs that we have developed, MyGene.info, MyVariant.info and MyChem.info. JSON-LD provides a standard way to add semantic context to the existing JSON data structure, for the purpose of enhancing the interoperability between APIs. We demonstrated several use cases that were facilitated by semantic annotations using JSON-LD, including simpler and more precise query capabilities as well as API cross-linking.

**Conclusions:** We believe that this pattern offers a generalizable solution for interoperability of APIs in the life sciences.

## Background

Recent developments in biological research have yielded a flood of data and knowledge about various biological entities, e.g. diseases, drugs, genes, variants, proteins and pathways. One key challenge that the scientific community faces is the large-scale integration of annotation information for each biological entity type, which is often fragmented across multiple databases. For example, ClinVar [1], dbSNP [2], and CADD [3] all contain useful and distinct information on human genetic variants. When filtering variants identified in a genome sequencing study, for example, large-scale integration of these variant data would greatly improve the efficiency of the data analysis. Moreover, a more important and sophisticated challenge would be to link data across multiple biological entities. For example, fields like systems chemical biology, which studies the effect of drugs on the whole biological system, requires the integration and cross-linking of data across from multiple domains, including genes, pathways, drugs as well as diseases [4].

Javascript Object Notation (JSON) is designed to be a lightweight, language-independent data interchange format that is easy to parse and generate. Over the past decade, JSON has been widely adapted in RESTful APIs. Previously, we have developed three biomedical APIs, MyGene.info [5, 6], MyVariant.info [7] and MyChem.info [8], that integrate public resources for annotations of genes, genetic variants and chemicals respectively. These three services collectively serve over six million requests every month from thousands of unique users. The underlying design and implementation of these systems are not specific to genes, variants or chemicals, but rather can be easily adapted to develop other entity-specific APIs for other biomedical data types such as diseases, pathways, species, genomes, domains and interactions.

A key challenge of Web API development is the semantic interoperability of data exposed by APIs. This challenge comes from the heterogeneity of biological entities (from variants to diseases), the variety of their data models and the scarcity of explicit data descriptions. Precise semantic alignment of individual JSON-based APIs would enable powerful integrative queries that span multiple web services, entity types, and biological domains.

JSON for Linking Data (JSON-LD) is currently being developed as a standard to promote interoperability among JSON-based web services [9]. JSON-LD offers a simple method to express semantically-precise Linked Data in JSON. It has been designed to be simple to implement, concise, backward-compatible, and human readable. JSON-LD is an official World Wide Web Consortium standard that has been well accepted and adopted, especially within the Internet of Things community [10]. While JSON-based web services are quite common among biomedical resources, the use of JSON-LD has not yet been extensively explored.

Here, we present our implementation and application of JSON-LD technology to biomedical APIs. We first implemented JSON-LD into three of the BioThings APIs we have developed, MyGene.info, MyVariant.info and MyChem.info. We then demonstrated its application by showing how it could be utilized to make data-structure neutral queries, to perform data discrepancy checks, to cross-link BioThings APIs as well as to integrate BioThings APIs into the linked data cloud. We believe that this work describes a generalizable pattern for stitching together individual APIs into a network of linked web services.

## Results

### Adding semantics to JSON document

JSON was specifically designed as a lightweight, language-independent data interchange format that is easy to parse and generate. However, the convenience and simplicity of JSON comes at a price, one of which is lack of namespace or semantics support.

Consider a scenario in which two biological data providers both created a key called “accession number” in their JSON document. One group used it to refer to a UniProt [11] accession number, while another group used it to refer to a ClinVar accession number. While a knowledgeable scientist can usually determine the intended usage, defining the semantics programmatically and automatically is much more difficult. It would be extremely useful for JSON documents to be "self-describing" in the sense that providers can explicitly define the semantic meaning of each key.

For the BioThings APIs, we solved this issue by implementing JSON-LD. Each API specifies a JSON-LD context (Figure 1), which itself is a JSON document and can provide a Universal Resource Identifier (URI) mapping for each key in the output JSON document. The use of URIs provides consistency when specifying subjects and objects. In our implementation, we used identifiers.org as the default URI repository. Identifiers.org [12] focuses on providing URIs for the scientific resources, especially in the Life Science domain. For example, a key for “rsid” could be assigned to the URI (“http://identifiers.org/dbsnp/”). Accessing this URI via HTTP shows a page with more detailed information about rsids. Importantly, a data consumer can be confident that two APIs that reference the same URI are referring to the exact same concept.

**Figure 1.**
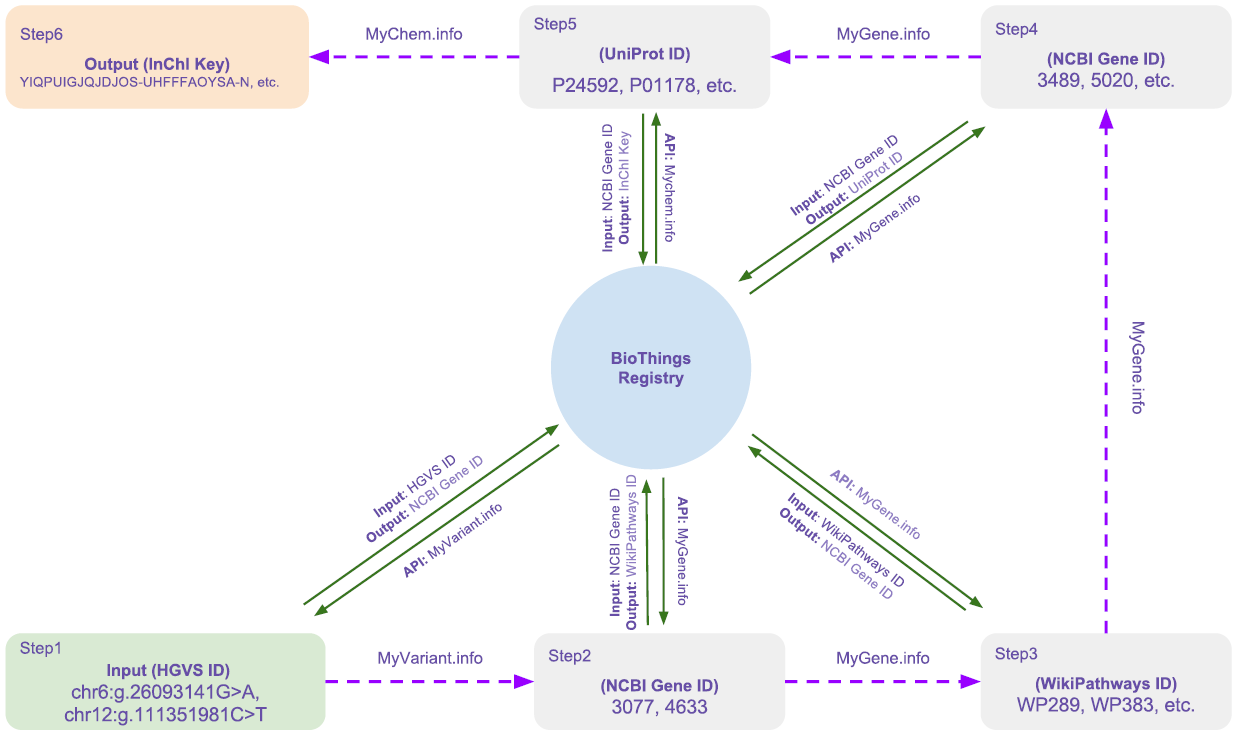
A simplified version of JSON-LD context for MyVariant.info. In this JSON-LD context, selected keys from the response document of MyVariant.info API are mapped to identifiers.org URIs. The complete version can be found at: http://myvariant.info/context/context.json

### Making data-structure neutral queries by URI

By using JSON-LD to define semantics in JSON documents, we found that JSON-LD could standardize and simplify the way we make queries in RESTful APIs. Most APIs are developed and maintained independently by different groups. And in most cases, API developers use different data representations and query syntax. Thus, a user has to read the API documentation and figure out the data structure and query syntax every time they need to handle a new API, which can be very time-consuming. For example, a user who would like to fetch the linked OMIM [13] disease IDs for a specific variant in MyVariant.info must first consult the JSON data schema, which would define the JSON field path of the OMIM ID (“clinvar.rcv.conditions.identifiers.omim”). Moreover, as services evolve, API developers often introduce incompatible changes in data structure between different versions, which would require API users to update their client code in order to properly handle the new JSON schema.

In contrast, an API that provides a JSON-LD context can be queried based on concept URIs in a way that is completely independent of the JSON data structure. Therefore, we adapted our *biothings_client* Python client [14] to use the JSON-LD context to translate data-structure neutral URIs into specific field locations within the JSON document [14]. In the example above, a user can simply query by the URI for OMIM ID (“http://identifiers.org/omim/”) without necessarily having to know the data source (ClinVar) and object structure. This pattern is demonstrated in **Additional file 1**. Although BioThings APIs are used in this demonstration, the JSON-LD processing procedure can be generalized to any JSON-based API that provides a JSON-LD context.

### Data discrepancy check

Variant annotation is a crucial part of the next genome sequencing data analysis. Incorrect annotations can cause researchers both to overlook potentially pathogenic DNA variants and to dilute interesting variants in a pool of false positives. Looking for discrepancies in data between different resources can be one way to assess data quality.

A typical data discrepancy check procedure would first requires the user to identify which data sources contain common data fields or identifiers (e.g. that dbNSFP and dbSNP both contain information about rsid). In addition, the user has to understand the query syntax and data structure of MyVariant.info in order to retrieve the data field from each data resource. JSON-LD, on the other hand, would greatly simplify the process by providing the semantic context of each data keys. As shown in Additional file 2, we performed a data discrepancy check based only on the JSON-LD context file. We were able to quantify the extent of differences in annotation of over 424 million variants from multiple variant annotation databases, including ClinVar (2017-04 release), dbSNP (version 150), and dbNSFP (version 3.4a) [15]. Of the 424 million variants in MyVariant.info, we found 10,842 unique variants that had different rsids reported across the resources that were imported (Additional file 3). Because each rsid can be unambiguously mapped to a unique genome location, this analysis clearly reveals some database-specific discrepancies (Additional file 4).

In addition to quality control, JSON-LD can also be utilized to conduct discovery-oriented queries. For example, we queried for variants with a high degree of variability in allele frequency in African populations as reported by 1000 genomes [16], ESP [17], and ExAC [18], a query that was greatly simplified by normalization to a common URI. We found 84 variants for which the reported allele frequency varied by more than 50% (Additional file 5), a list that could be be notable for studying different selective pressures among African sub-populations.

### Facilitate API Cross Linking

When JSON-LD contexts are used, annotating data between genes, variants and drugs can be seamlessly integrated together. For example, consider a use case where an upstream analysis identified two missense variants (“chr6:g.26093141G>A” and “chr12:g.111351981C>T” in hg19 genome assembly) related to a rare Mendelian disease. If an analyst wanted to obtain genes which the variant might affect as well as available drugs targeting these affected genes, a typical workflow would involve the following steps (Figure 2):

a. Query MyVariant.info to retrieve the annotation objects for variant “chr6:g.26093141G>A” and “chr12:g.111351981C>T”
b. Parse the annotation objects to get the NCBI Gene IDs related to these variants from the dbsnp.gene.geneid field, which are “3077” and “4633”
c. Query MyGene.info to retrieve the annotation objects for NCBI Gene “3077” and “4633”
d. Parse the annotation objects to get the WikiPathways [19] IDs related to each gene from the pathway.wikipathways.id field
e. Query MyGene.info again to retrieve annotation objects related to the WikiPathways IDs from step d.
f. Parse the query results to retrieve all UniProt IDs related to these WikiPathways IDs.
g. Query MyChem.info to retrieve the drug objects associated to the target proteins, using the UniProt IDs found in step f.

**Figure 2.**
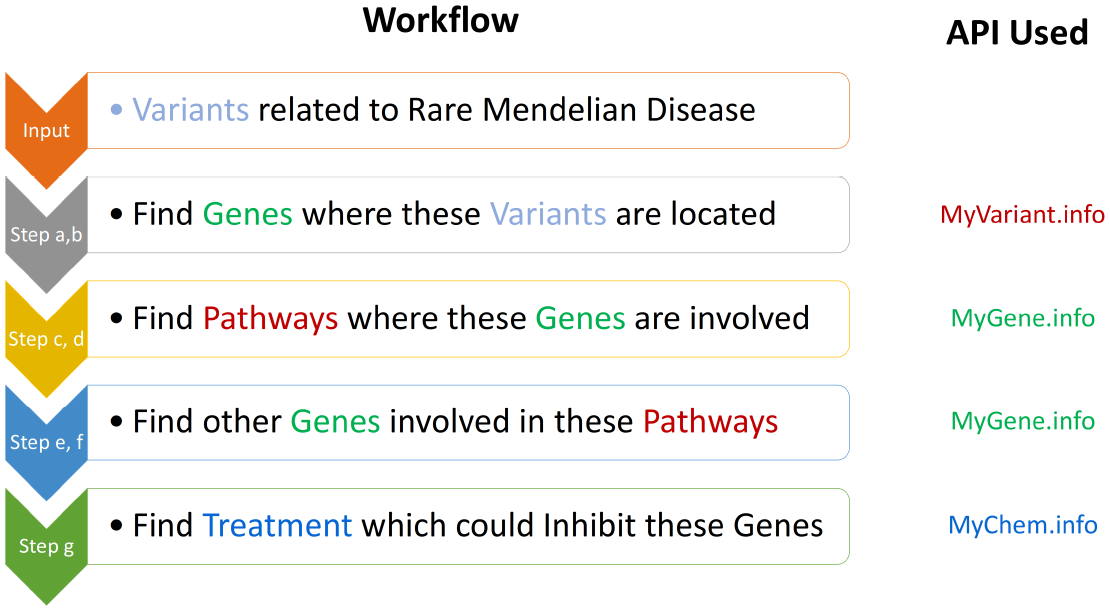
Typical workflow for finding treatments related to a rare Mendelian disease using multiple BioThings APIs. An upstream analysis first identified some variants related to a rare Mendelian disease. Next, the analyst want information about genes where these variants are located (step a, b). Then, from the genes, the analyst would like to know all the pathway information where these genes are involved (step c, d). Furthermore, the analyst would also like to know other genes involved in these pathways (step e, f). Finally, the analyst want to obtain information about all available treatment options (e.g. drugs) available targeting all these genes obtained in the previous steps (step g).

To perform this workflow, traditionally, users must first manually inspect the output or the documentation of MyGene.info, MyVariant.info and MyChem.info to identify the relevant JSON keys (e.g. dbsnp.gene.geneid and pathway.wikipathways.id).

An alternative approach utilizing JSON-LD greatly simplified the protocol (Figure 3). We first created a BioThings API Registry that was constructed from the JSON-LD context files of all BioThings APIs [20]. The registry records information about the input and output type for each API (expressed as URIs) as well as the query syntax for each endpoint of BioThings APIs (as shown in Additional file 6). This registry was then used to help locate the right API to use when input/output type is specified. JSON-LD, on the other hand, could be utilized to construct the API query and extract the output from the JSON document. As demonstrated in Additional file 7, users no longer need to have any prior knowledge of API input and output data structures. It only requires the user to specify the URI for the input type (e.g. http://identifiers.org/ncbigene/) and the target output type (e.g. http://identifiers.org/wikipathways/). We implemented a Python function *IdListHandler*, as shown in the demo code, which automatically scans through the JSON-LD contexts provided from a list of APIs, currently demonstrated by our existing BioThings API (MyGene.info, MyVariant.info and MyChem.info). *IdListHandler* then selects the API which is able to perform the task and automatically executes the queries to return the desired output. As we are expanding the scope of BioThings API to cover additional biomedical entities (e.g. diseases, phenotypes), the power of this approach will continue to grow. Moreover, since the JSON-LD context can be provided by a third-party, not just the API providers, the mechanism we described here can be further extended to cross-link even broader scope of biomedical APIs.

**Figure 3.**
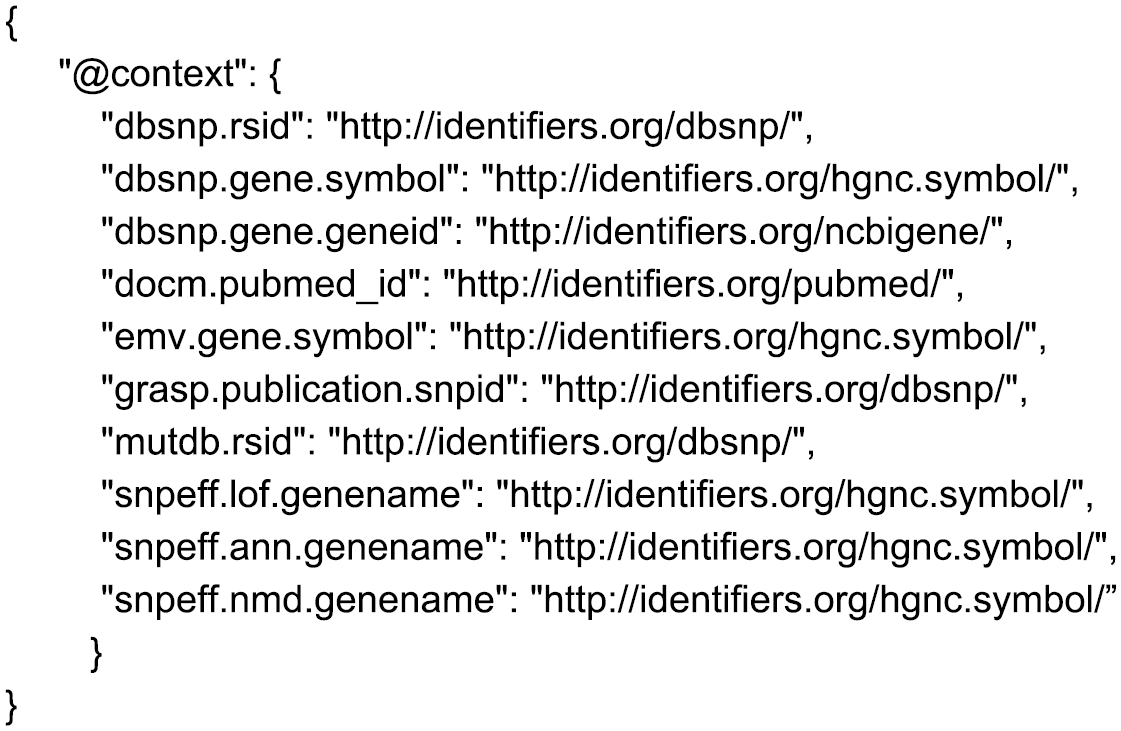
Workflow of using JSON-LD and BioThings API Registry to perform complex queries. By combining JSON-LD and BioThings API Registry, the Python function *IdListHandler* is able to select the API which is able to perform the task and automatically executes the queries to return the desired output. Users could easily get a list of drug candidates related to a rare Mendelian disease by the following path: HGVS ID → NCBI Gene ID → WikiPathways ID→ NCBI Gene ID → UniProt ID → Drug InChI Key. It only requires the user to specify the input/output type at each step.

### Easy conversion to RDF

The Resource Description Framework (RDF) has been widely used to describe, publish and link data in the life science domain. It generates a series of "triples" consisting of a subject, predicate and object.

A number of biomedical resources provide RDF to facilitate relationship exploration, such as Bio2RDF [21], Monarch [22], and Open PHACTS [23]. Since JSON-LD is simply a JSON-based representation of Linked Data, it is programmatically simple to export JSON-LD data into RDF for integration with other RDF-based resources (demonstrated in Additional file 8). JSON-LD offers a convenient method by which bioinformatics web services (like MyGene.info, MyVariant.info and MyChem.info) can be integrated with the network of Linked Data.

## Discussion

The biomedical research community has seen a proliferation of web services in recent years. These services have become a primary route for the bioinformatics community to disseminate and consume data and analysis methods. Individually, these web services are very useful components of our informatics ecosystem. Nevertheless, there is growing appreciation that creating an integrated network of APIs would be an even more powerful resource that is greater than the sum of its parts.

Here, we demonstrate that JSON-LD is one technical solution for integration of web-based APIs. JSON-LD offers many advantages – that it builds on the widely-used JSON data exchange format, that it itself is a W3C standard, and that it offers the potential of decoupling the authoring of semantic context from the serving of data. In fact, several biomedical tool providers have already introduced JSON-LD in their APIs, including Monarch, CEDAR [24], and UniProt.

Nevertheless, two key challenges remain before achieving broader adoption and the ability to address more complex biomedical use cases. First, there needs to be greater standardization of URIs for biological concepts. In addition to identifiers.org, other entities like health-lifesci.schema.org [25] and Bio2RDF provide URIs for biologically-relevant entities. Second, a mechanism for expressing in a structured format the semantic nature between concepts and the provenance of such relationships would expand the richness of possible queries in a JSON-LD ecosystem.

## Conclusions

We believe that the proofs-of-concept presented here demonstrate that the JSON-LD pattern described here already have useful applications, and that adoption of this pattern would greatly expand the interoperability of biomedical web services.

## Methods

### URI Repository

In our implementation, we used http://identifiers.org as the default URI repository, which provides URIs for the scientific community.

### Creating JSON-LD Context

JSON-LD context for each BioThings API is created by mapping individual field name to URI. For example, a field named "dbsnp.rsid" in MyVariant.info is mapped to http://identifiers.org/dbsnp/. Example BioThings API context could be found at http://myvariant.info/context/context.json

### API Cross-Linking

API cross-linking is made possible through the combination of BioThings API Registry and JSON-LD transformation.

BioThings API Registry records information about the input and output type accepted as well as the query syntax for each endpoint of BioThings APIs. It is utilized to find the right API given input and output type specified by the user.

JSON-LD transformation is performed using PyLD python package (version 0.7.2) [26]. When a JSON document is retrieved along with its JSON-LD context, JSON-LD transformation will convert the JSON document into N-Quads format where each value is mapped to an URI. Through N-Quads output, we can then extract the desired output data.

### RDF Transformation

RDF transformation is performed using PyLD python package (version 0.7.2) [26]. It is demonstrated using Jupyter Notebook stored in github (S8 Code Example).

## Abbreviations

API: Application Programming Interfaces
JSON: Javascript Object Notation
RDF: Resource Description Framework
URI: Universal Resource Identifier
JSON-LD: JSON for Linked Data

## Declarations

### Ethics approval and consent to participate

Not applicable.

### Consent for publication

Not applicable.

### Availability of data and materials

The information for data, workflows and scripts used in the paper is available under an Apache software license at https://github.com/biothings/JSON-LD_BioThings_API_DEMO

## Authors’ contributions

Conceived and designed the methodology: AIS CW JX. Performed the implementation and data analysis: JX SL CA JA. Wrote the paper: JX. Work supervised by AIS CW. All authors read and approved the final manuscript.

## Supporting Information

1. **Additional file 1. A Jupyter Notebook demonstration of how to Make Data-structure Neutral Queries by URI.**
2. **Additional file 2. A Jupyter Notebook demonstration of how to perform data discrepancy check using JSON-LD**
3. **Additional file 3. Table listing 10,842 unique variants that had different rsids reported across the resources recorded in MyVariant.info.**
4. **Additional file 4. Figure showing an example of data discrepancy for rsid between different sources.** Data discrepancy for rsid between different sources, e.g. dbNSFP, dbSNP and mutDB. The default assembly used in dbNSFP is hg38. And the hg19 genomic positions provided by dbNSFP were lifted over from its hg38 positions, which is corresponding to a different rsid (“rs542852754”) in this case from the one dbSNP provides (“rs7182058”). This suggests an error caused in the liftover process.
5. **Additional file 5. Table listing 84 variants for which the reported allele frequency varied by more than 50% across the resources recorded in MyVariant.info.**
6. **Additional file 6. A Jupyter Notebook demonstration of BioThings API registry.**
7. **Additional file 7. A Jupyter Notebook demonstration of how to cross-link BioThings APIs in order to perform complex queries across APIs.**
8. **Additional file 8. A Jupyter Notebook demonstration of how to convert a JSON document from MyVariant.info into RDF using JSON-LD.**

